# Integrated Molecular Characterization of Intraductal Papillary Mucinous Neoplasms: An NCI Cancer Moonshot Precancer Atlas Pilot Project

**DOI:** 10.1101/2022.10.14.512148

**Authors:** Alexander Semaan, Vincent Bernard, Justin Wong, Yuki Makino, Daniel B. Swartzlander, Kimal I. Rajapakshe, Jaewon J. Lee, Adam Officer, Christian Max Schmidt, Howard H. Wu, Courtney L. Scaife, Kajsa E. Affolter, Daniela Nachmanson, Matthew A. Firpo, Michele Yip-Schneider, Andrew M. Lowy, Olivier Harismendy, Subrata Sen, Anirban Maitra, Yasminka A. Jakubek, Paola A. Guerrero

**Author notes:** equal contribution (Alexander Semaan, Vincent Bernard).

## Abstract

Intraductal papillary mucinous neoplasms (IPMNs) are cystic precursor lesions to pancreatic ductal adenocarcinoma (PDAC). IPMNs undergo multistep progression from low grade (LG) to high grade (HG) dysplasia, culminating in invasive neoplasia. While patterns of IPMN progression have been analyzed using multi-region sequencing for somatic mutations, there is no integrated assessment of molecular events, including copy number alterations (CNAs) and transcriptomics changes, that accompany IPMN progression. We performed laser capture microdissection on surgically resected IPMNs of varying grades of histological dysplasia obtained from 24 patients (total of 74 independent histological lesions), followed by whole exome and whole transcriptome sequencing. Overall, HG IPMNs displayed a significantly greater aneuploidy score than LG lesions, with chromosome 1q amplification, in particular, being associated with HG progression and with cases that harbored cooccurring PDAC. Furthermore, the combined assessment of single nucleotide variants (SNVs) and CNAs identified both linear and branched evolutionary trajectories, underscoring the heterogeneity in the progression of LG lesions to HG and PDAC. At the transcriptome level, upregulation of MYC-regulated targets and downregulation of transcripts associated with the MHC class I antigen presentation machinery was a common feature of progression to HG. Taken together, this work emphasizes the role of 1q copy number amplification as a putative biomarker of high-risk IPMNs, underscores the importance of immune evasion even in non-invasive precursor lesions, and supports a previously underappreciated role of CNA-driven branching evolution as an avenue for IPMN progression. Our study provides important molecular context for risk stratification and cancer interception opportunities in IPMNs.

**Significance:** Integrated molecular analysis of genomic and transcriptomic alterations in the multistep progression of intraductal papillary mucinous neoplasms (IPMNs), which are bona fide precursors of pancreatic cancer, identifies features associated with progression of low-risk lesions to high-risk lesions and cancer, which might enable patient stratification and cancer interception strategies.

## Introduction

Pancreatic ductal adenocarcinoma (PDAC) remains a disease with a dismal prognosis and is estimated to become the second leading cause of cancer related death in the United States within the next decade [1, 2]. Although most patients with PDAC present with locally advanced or metastatic disease, a minority of patients are diagnosed with localized disease where curative resection remains an option. Detection and intervention of disease at a localized stage results in a significant survival benefit. PDAC is thought to arise from two distinct subtypes of precursor lesions – approximately 85-90% of cancers occur on the backdrop of microscopic precursor lesions known as pancreatic intraepithelial neoplasia or PanIN. The remaining ~10-15% are believed to arise from mucinous cystic precursor lesions, of which the vast majority are intraductal papillary mucinous neoplasms (IPMNs). Given the projected timeline of several years over which non-invasive precursor lesions progress to invasive neoplasia [3], there is a potentially wide window of opportunity for early detection of this lethal neoplasm.

While PanINs are typically not amenable to non-invasive imaging-based detection, IPMNs have the benefit of being detectable on conventional abdominal imaging studies. Nonetheless, the currently used clinical algorithms, while representing a considerable improvement in stratifying which patients should merit surgery compared to prior schema, continue to both miss incident cancers and overestimate cancer risk in cysts that can be managed conservatively [4]. Elucidating the molecular underpinnings of IPMN progression could provide an avenue for identifying those lesions which might be at greatest risk and generate opportunities for early cancer interception.

The genomic landscape of IPMNs has been steadily cataloged over the past decade, and has identified both early drivers that predominate in LG IPMNs, such as *KRAS, GNAS* and *RNF43* mutations, as well as drivers associated with IPMN progression, including *TP53, PIK3CA* and *SMAD4*, amongst others [5]. Recent studies have also revealed genomic heterogeneity within different regions of IPMNs, suggesting most IPMNs originate as polyclonal lesions prior to emergence of a dominant clone [6, 7]. Nonetheless, an integrated molecular analysis of IPMNs of varying histological grades that combines global genomic-wide and transcriptomic analyses has been lacking. This approach can provide unique insights into non-genomic mechanisms of cellular perturbation driving IPMN progression, including potential cross talk mechanisms with the precursor microenvironment (PME).

Our study was performed as part of the NCI Cancer Moonshot Precancer Atlas Pilot Project (PCAPP), which is a component of the publicly funded NCI Human Tumor Atlas Network (HTAN) [8]. In this study, we performed integrated whole exome and transcriptomic sequencing, of LG and HG IPMNs. Our cohort included both independent HG IPMNs (and PDAC), as well as synchronous HG IPMNs arising in the context of a pre-existing LG neoplasm. The latter subset is uniquely informative in identifying whether molecular aberrations seen in HG lesions are “wired in” at an earlier stage of dysplasia. We were able to validate many of the “early” and “late” genomic drivers previously reported in IPMN pathogenesis, but also elucidated previously unreported CNAs that stratified LG IPMNs at risk of progression to HG IPMNs and PDAC. Additionally, we identify that predominantly HG lesions with an invasive component do not follow a linear evolution, with CNAs branching driving evolution of subclones. At the transcriptomic level, downregulation of transcripts related to antigen presentation was a pervasive feature of IPMN progression, establishing that immune evasion reported in PDAC has its origins in non-invasive lesions.

## Methods

### Patient cohort

The HTAN PCAPP in various precursor lesions was organized under the umbrella of the NCI-funded Consortium for Molecular and Cellular Characterization of Screen-Detected Lesions Create (MCL) (https://mcl.nci.nih.gov), with MD Anderson leading the PDAC precursor atlas effort. For this PCAPP, a total of 74 histological samples of IPMN cystic lesions and 10 blood samples from 24 patients were collected between March 2016 and February 2019 across the United States: At the MD Anderson Cancer Center (MDACC) under protocols Lab00-396, PA11-0670, at the Indiana University (IU) under protocol number 1011003217 (0209-66), at the University of California San Diego (UCSD) under protocol number IRB_151608 and at the University of Utah (UU) under protocol numbers IRB_00011467 and IRB_00089989. All but two patients had normal tissue or PBMC for germline correction (10 peripheral blood mononuclear cells (PBMCs), 9 normal pancreatic tissue, two splenic and one duodenal tissue). The study was performed in accordance with standard ethical guidelines approved by the institutional review board (IRB) at every site and in accordance with the Declaration of Helsinki. All patients had clinically and histologically confirmed intrapapillary mucinous neoplasm (IPMN). In compliance with the PCAPP guidelines, formalin-fixed paraffin-embedded (FFPE) blocks with a median of 16 months after resection were used (range 2 - 33 months). There was no correlation between block age and DNA integrity across institutions which might indicate differences in tissue processing between centers (R^2^=0.13, p=0.08, **Supplementary Figure 1A**).

### Laser microdissection (LCM), isolation and quality control

All FFPE slides were reviewed by an experienced pancreas/GI pathologist at the contributing site and verified at MDACC by one of the authors and an expert pancreas pathologist (A.M.). The histological grade was assigned in accordance with the updated guidelines for pre-neoplastic precursor lesions in the pancreas [9]. LCM and library preparations were centrally performed at MDACC for all samples. Depending on availability, as many as five different areas per patient were collected via LCM using the PALM MicroBeam system (Carl Zeiss Microscopy GmbH, Jena, Germany) for both DNA and RNA isolation. These area types include normal ducts (ND), acinar cells (AC), low-grade IPMN (LG) and high-grade IPMN (HG) lesions, and PDAC.

An average of three (range 1-7 slides) consecutive 7μm, hematoxylin eosin (HE)-stained FFPE slides were used for DNA extraction and pooling from each compartment (ND, AC, LG or HG) and a median of three compartments was dissected per patient (range 1-4). DNA isolation was performed using the QIamp DNA Micro Kit (Qiagen, Hilden, Germany cat # 56304) with a modified protocol: 18μl of buffer ATL and 12μl of Proteinase K were combined by vortexing and applied to a customized AdhesiveCap 200 clear (D) (Carl Zeiss Microscopy GmbH, Jena, Germany, cat # 415190-9191-000). Samples were then incubated overnight in an upside-down position at 56 °C. Twenty-five microliter of buffer ATL and 50μl of buffer AL were added and pulse-vortexed for 10 sec. Following, 50μl of ethanol (100%) were added, pulse-vortexed for 10 sec and incubated for 5 min at room temperature. Lysate was then transferred to QIAamp MinElute columns and centrifuged at 6000 x g for 1 min. Subsequently, 2 washing steps at 6000 x g for 1 min with 500μl of buffer AW1 and 500μl of buffer AW2 were followed by a drying step (20,000 x g for 3 min). Columns were incubated with 20μl of distilled deionized water for 10 min and finally centrifuged at 20,000 x g for 1 min to elute DNA. The DNA obtained from each compartment was then pooled and volume was reduced using the Savant™ SpeedVac™ DNA 130 Integrated Vacuum Concentrator System (Thermo Fisher, Waltham, USA, cat # DNA130-115). Pooled DNA concentration was measured using the Qubit™ dsDNA BR Assay Kit (cat #: Q32853, Qubit 2.0 fluorometer). DNA was stored at −20C until further processing.

Additionally, bulk germline DNA (gDNA) was extracted from two 7um FFPE using the QIAamp DNA FFPE Tissue Kit (Qiagen, Hilden (Germany), cat # 56404). gDNA integrity was measured by the genomic DNA ScreenTape (Agilent, Santa Clara, cat # 5067-5365) on a Tapestation 2200 system in conjunction with TapeStation Analysis Software (Agilent, Santa Clara). Median DNA Integrity Number (DIN) was 4.95 (range 2.4 – 6.2). Matching whole blood samples were collected in acid citrate dextrose tubes (BD) and processed within 3-4 hours of phlebotomy (n=10) as previously described [10]. Whole blood was centrifuged at 2500 x g for 10 mins to separate plasma. PBMCs were isolated using the Lymphocyte Separation Medium (Corning, cat # 25-072-CV) and centrifugation at 620 x g for 30 mins. PBMC DNA was isolated using the DNeasy Blood & Tissue Kit (Qiagen, Hilden, cat # 69506) following the manufacturer’s protocol.

### DNA Library construction and sequencing

A median of 63ng (range 10-200ng) for FFPE derived, pooled DNA and a median of 155ng (range 105-200ng) of matched-PBMC DNA was fragmented using the SureSelect XT HS and XT Low Input Enzymatic Fragmentation Kit following the manufacturer’s instructions (Agilent, Santa Clara, USA, cat # 5191-4080). Molecular-barcoded libraries were constructed following the SureSelect XT HT targeted enrichment protocol for Illumina paired-end multiplexed sequencing libraries (Version A1, July 2017) as previously described [10] with the following modifications: Step 2.4: Incubation at 20°C for ligation for 35 min, step 3.1: Pre-capture pooling of samples were used if necessary to reach a minimum input for the hybridization step of 50ng and incubation temperature for segment numbers 2-5 were reduced to 62.5°C. SureSelect Clinical Research Exome V2 (Agilent, Santa Clara, USA, catalog # 5190-9492) and Exome V7 (Agilent, Santa Clara, USA, catalog # 5191-4005) was used for capturing. For cross validation samples (n=6) the All-In-One solid tumor panel (AIO, Agilent, Santa Clara, USA, catalog # G9706S) was used for capture. Final libraries were multiplexed, denatured and diluted to a final concentration of 1.7 pM for sequencing and cluster generation as per manufacturer’s recommendation. Clustered flow-cells were sequenced on the Illumina NextSeq 500 instrument targeting 400x coverage (Illumina, San Diego, USA) using standard Illumina paired-end primers and chemistry (Index 1=8, index2=10, read length=125).

### Analysis of mutations and copy number alterations (CNA)

#### Alignment and processing

Raw sequencing data were converted to fastqs with bcl2fastq (v2.20.0.422), including a molecular barcode fastq. Fastq files were assessed for quality with FastQC (v0.11.8), trimmed with SureCall Trimmer (AGeNT, Agilent, v4.0.1) to remove adaptor sequences, and then aligned to hg19 with BWA (0.7.15-r1140). The resulting BAM was then collapsed by barcodes to family size of one, according to default parameters for LocatIt (AGeNT, Agilent, v4.0.1) with two exceptions: without filtering for barcode quality (q=0), and correcting for optical duplicate detection (c=2500) to account for sequencing on patterned flow cells. Collapsed bams were then processed for base quality score recalibration (BQSR) according to best practices by the genome analysis toolkit (GATK, 4.1.2.0), using dbSNP138 to exclude consideration. Samples that failed quality control (QC) were re-sequenced.

#### Mutation Calling

Three callers were used for somatic variant detection: Mutect2 (GATK, v4.1.3.0), SureCall (Agilent, 4.1.1.9), and MuSE (v1.0rc). Across all three callers, tumor samples were run against a paired normal based on availability (order priority: PBMC, uninvolved non-pancreatic tissue, normal ducts (ND)). Mutect2 was run according to best practices and default parameters, including checking for cross-sample contamination, and filtering for sequencing artifacts (such as orientation-based FFPE artifacts). To further reduce possible population variation, a panel of normal DNA with all available PBMCs and NDs (without detectable *KRAS* and *GNAS* mutations) was used as an additional filter. When no paired normal was available, Mutect2 was run in tumor-only mode, filtering against the panel of pooled normal sequences. A set of high-confidence Mutect2 calls was generated by further filtering with a CONTQ score of 50 or greater. SureCall was run from BAMs according to the Agilent-provided defaults: “Default SureSelect Tumor Normal Method” when paired data was available, or by “Default SureSelect” method in single sample mode when not. MuSE was also run according to default parameters, and variants meeting the “PASS” or “Tier1” criteria were included in downstream analyses. Since matched normals are required for MuSE, two samples (5 and 19) did not have MuSE calls. Lastly, VCFs were annotated by ANNOVAR (2018-04-16) across a variety of metrics, including gnomAD, ExAC (non-TCGA), COSMIC86, and ClinVar. SNVs called by two or more of the callers were used for downstream analysis. When MuSE calls were unavailable, the union of Mutect2 and SureCall SNV calls were used for downstream analysis. All SNVs with an allele frequency greater than 1% in gnomAD or ExAC (non-TCGA), were removed.

The SNV call set from this agnostic approach was complemented by a set of variants identified through a targeted analysis at pre-determined genomic locations identified by only one caller and requiring less stringent filtering parameters. These predetermined genomic loci included two groups, those in established PDAC driver genes and SNVs detected by the more stringent approach in other samples from the same patient. These included a set of mutect2 variant calls that were generated under more sensitive conditions (lowering the log odds threshold for emission to 1.5). PDAC driver genomic loci were defined as those with 15 or more entries in the COSMIC database in established PDAC genes [11, 12] (*KRAS; TP53; SMAD4; CDKN2A; GNAS; BRAF; PIK3CA; MAP2K4; TGFBR1; TGFBR2; RNF43; CTNNB1; STK11; ARID1A; KDM6A; SF3B1; RBM10; IDH1; PTEN; APC; ATM; BRCA1; BRCA2*). Additionally, SNVs were called using this targeted approach at genomic sites where an SNV had been detected using the two-caller approach in at least one of the patient’s other non-blood samples.

SNVs were classified as deleterious if they were an exonic or splicing variant, and if they were labeled as deleterious by two or more prediction models (sift, polyphen, HVAR, LRT, mutationTaster, fathmm, provean), or if the variant was labeled as pathogenic in ClinVar. Small insertion and deletion calls were generated using Pindel (0.2.5b9) with the default parameters and filtered by depth (minimum 5 supporting reads in tumor, maximum 0 corresponding reads in normal). Analysis of mutation burden and detection of CNAs was performed as previously described [10].

To ensure high quality SNV calling, we performed extensive cross validation including parallel ddPCR mutation calling for *KRAS* and *GNAS* in all samples and parallel, ultra-deep targeted panel sequencing. Both methods showed a high concordance of the called mutations (ddPCR: R^2^=0,95, p<0.0001), **Supplementary Table 1, Supplementary Figure 1B**.

### TCGA 1q analysis

Previously published TCGA PDAC data were analyzed in order to determine an association between the RNA molecular subtype and 1q whole arm amplification. RNA based classifications (Moffitt, Collissson, and Bailey) and copy number calls were obtained from previous publications [11, 13]. This analysis was limited to TCGA PDAC samples with a consensus classification of basal or classical for all three classifiers. A chi-square test was used to determine significance (18 classical with 1q gain, 16 classical without 1q gain, 3 basal with 1q gain, and 15 basal without 1q gain; p = 0.017).

### RNA isolation, Library construction and Sequencing

For RNA, tissues were harvested directly into caps by LCM as described above, collecting 500-1000 cells as input per library, and stored at −80C until ready for use. We processed the tissue via a modified SMART3 protocol [14]. Briefly, we performed steps as described by Foley et al. up to PCR. PCR was performed as described except for cycle number (20, 21, or 22) was additionally dependent on planned post-PCR replicate pooling (3, 2, or 1 respectively). Following PCR amplification, libraries from the same patient and tissue were pooled, cleaned up using beads, and stored in RNase-free water. Samples were indexed with indices 1-16, 18-20, 22, 25, and 27 from, [14] corresponding to TruSeq LT indices. Prepared libraries were quantified and checked for appropriate size distribution using an RNA ScreenTape (Agilent, Santa Clara, cat # 5067-5579) on a Tapestation 2200 system in conjunction with TapeStation Analysis Software (Agilent, Santa Clara), and stored at [-80] until ready for sequencing. When ready, libraries were pooled, multiplexed, and diluted to a final concentration of 1.6 pM for sequencing on a NextSeq 500 (Illumina, San Diego, USA), single-end, with 75 cycles (Illumina, cat# 20024906).

Raw files were processed as described previously [14]. Briefly, raw data from the sequencer were converted to fastqs with bcl2fastq (v2.20.0.422) and fastqs were assessed for quality with FastQC (v0.11.8). The 3SEQtools suite (https://github.com/jwfoley/3SEQtools) was used to further process the data including read trimming for adaptor contamination and polyA tails (FastQC), alignment with STAR (2.7.1a), and depth-aware deduplication (3SEQtools). Finalized bams were processed for differential expression using DESeq2.

In order to infer PDAC molecular subtypes in our samples, we used an approach previously utilized in hepatocellular carcinonoma [15]. SCnorm was used to normalize expression as it has been used for scRNA-seq [16]. Subtype classification was done using nearest template prediction and the package CMScaller was used as a wrapper for the nearest template prediction function. Finally, we applied the Moffitt [17], Collison [18] and Bailey [19] classifiers to each RNAseq replicate.

### Digital Droplet PCR (ddPCR) analysis

Droplet digital polymerase chain reaction (ddPCR) was performed using whole-slide FFPE extracted DNA on a QX200 Droplet Digital PCR System (BioRad, Hercules USA) following previously described protocols [20, 21]. For highly sensitive multiplex *KRAS* and *GNAS* detection, we used specific *KRAS* probes following the manufacturer’s protocol (BioRad, Hercules USA) for G12V (cat. # dHsaMDV2510592), G12D (cat. # dHsaMDV2510596), and G12R (cat. # dHsaCP2506874) and GNAS probes (BioRad, Hercules USA) R201S (cat. # dHsaMDS2513808), R201C (cat. # dHsaMDV2510562) and R201H (cat. # dHsaMDV2516796).

### Neo-epitope prediction

Personalized human leukocyte antigen (HLA) types were generated from the WES data using OptiType (1.3.1) and confirmed using PolySolver (1.0). For Optitype, finalized DNA bams were deconstructed back into paired-end fastqs with bedtools (bamtofastq, 2.28.0). Each paired-end fastq was then processed separately and aligned to an HLA-reference with RazerS3 (3.5.7). The resulting BAMs are then deconstructed a second time with samtools (bam2fq, 1.9), to create fished/filtered fastqs. These fastqs are then processed together with OptiType with default parameters to generate an HLA-type for each person/tissue. For PolySolver, the finalized bams are processed directly. HLA-types are considered confirmed when both OptiType and PolySolver agree.

Personalized variant antigens by cancer sequencing (pVACseq), from the cancer immunotherapy tools suite, pVACtools (v1.5.2) was then used for neo-epitope prediction. VCFs from both the mutect2 and MuSE pipelines were filtered by the list of consensus calls, annotated with Variant Effect Predictor (VEP, v94), to which coverage was also added using kallisto (0.44.0). We also added expressions of the transcript according to the SMART3 data with the python package VAtools (vcf-readcount-annotator) and kallisto. Pindel calls were restricted to established cancer genes [22] and processed similarly. The VEP-, coverage- and expression-annotated VCFs are then processed with pVACseq, using the confirmed personalized HLA-types, an epitope size ranging from 8-11, and with multiple algorithms specified (MHCflurry, MHCnuggetsI, NetMHC, NetMHCpan, PickPocket, SMM, and SMMPMBEC). The resultant list of predicted neoepitopes was then combined per patient/tissue, checked for duplicates (e.g. variants called in both Mutect2 and MuSE).

### Statistical Analysis

Statistical analyses were performed with Prism 8 (Graph Sotfware, Inc.) and statistical significance was determined as a P value of <0.05.

## Results

### IPMN cohort and clinicopathological features

A total of 74 histological samples from 24 surgically resected IPMN cases were analyzed for their genome and transcriptome expression. One sample failed initial QC control and was furthermore excluded from analysis. Two of the processed FFPE blocks for analyses contained invasive carcinoma, although 9/24 of the final pathological reports indicated an invasive carcinoma (median maximum diameter of 6mm, range <1mm - 50mm). Patient data and clinical annotations are summarized in **Table 1.** and an experimental outline can be found in **Supplementary Figure 1C-D.**

**Table 1:** Patient demographics and clinicopathological characteristics.

| **Cohorts Characteristics** | **Patients** |
| --- | --- |
| **Total patient number** | 24 |
| **Peripheral blood samples** | 10 |
| **Non-tumorous tissue sample** | 9 pancreatic tissue, 1 duodenum, 2spleen |
| **Age of resection (median)** | 70 years (range 56-82 years) |
| **<u>Gender</u>** |  |
| **Men** | 15 (62.5%) |
| **Women** | 9 (37.5%) |
| **Maximum diameter IPMN (median)** | 27.5mm (range 7-50) |
| **Carcinoma detected** | 9 (37,5%) |
| **<u>Histological grading</u>** |  |
| **Intermediate** | 9 (37.5%) |
| **High** | 15 (62.5%) |
| **<u>Histological subtypes IPMN</u>** |  |
| **Intestinal** | 8 (33.3%) |
| **Pancreatobiliary** | 12 (50%) |
| **Gastric** | 4 (16.6%) |
| **<u>Relation to main duct</u>** |  |
| **Side branch** | 6 (25%) |
| **Main-duct** | 6 (25%) |
| **Main and side branch** | 3 (12,5%) |
| **Not reported** | 9 (37,5%) |
| **<u>Cyst location</u>** |  |
| **Head** | 13 (54.1%) |
| **Tail/Body** | 10 (41.7%) |
| **Diffused** | 1 (4.2%) |

### *KRAS* and *GNAS* mutations in IPMNs

Our whole exome sequencing approach indicated that most premalignant lesions either harbored *GNAS* or *KRAS* mutations (91.3% and 65.2%, respectively). Nine patients had the same *KRAS* and *GNAS* mutations in both their LG and HG samples. In addition, 2 patients harbored the same *KRAS* mutations in their LG and HG samples and 2 patients had the same *GNAS* mutations in their LG and HG samples (**Figure 1A**). These data support a model where majority of patients with LG and HG IPMNs have a shared mutation in KRAS and/or GNAS (i.e. having arisen in the same clonal lineage or a truncal mutation). For patient 10, we documented a shared *KRAS* mutation in the LG and ND supporting prior observations that these founder clones may be present in pathologically normal appearing tissues. In four patients (three LG to HG and one HG to PDAC), we detected multiple *KRAS* mutations in the LG (patient IDs 6, 19, and 24) and HG (patient ID 16) pointing to clones that independently acquired *KRAS* mutations, suggestive of strong convergent evolution for clones with *KRAS* mutations in pancreatic tissues from these patients.

**Figure 1:**
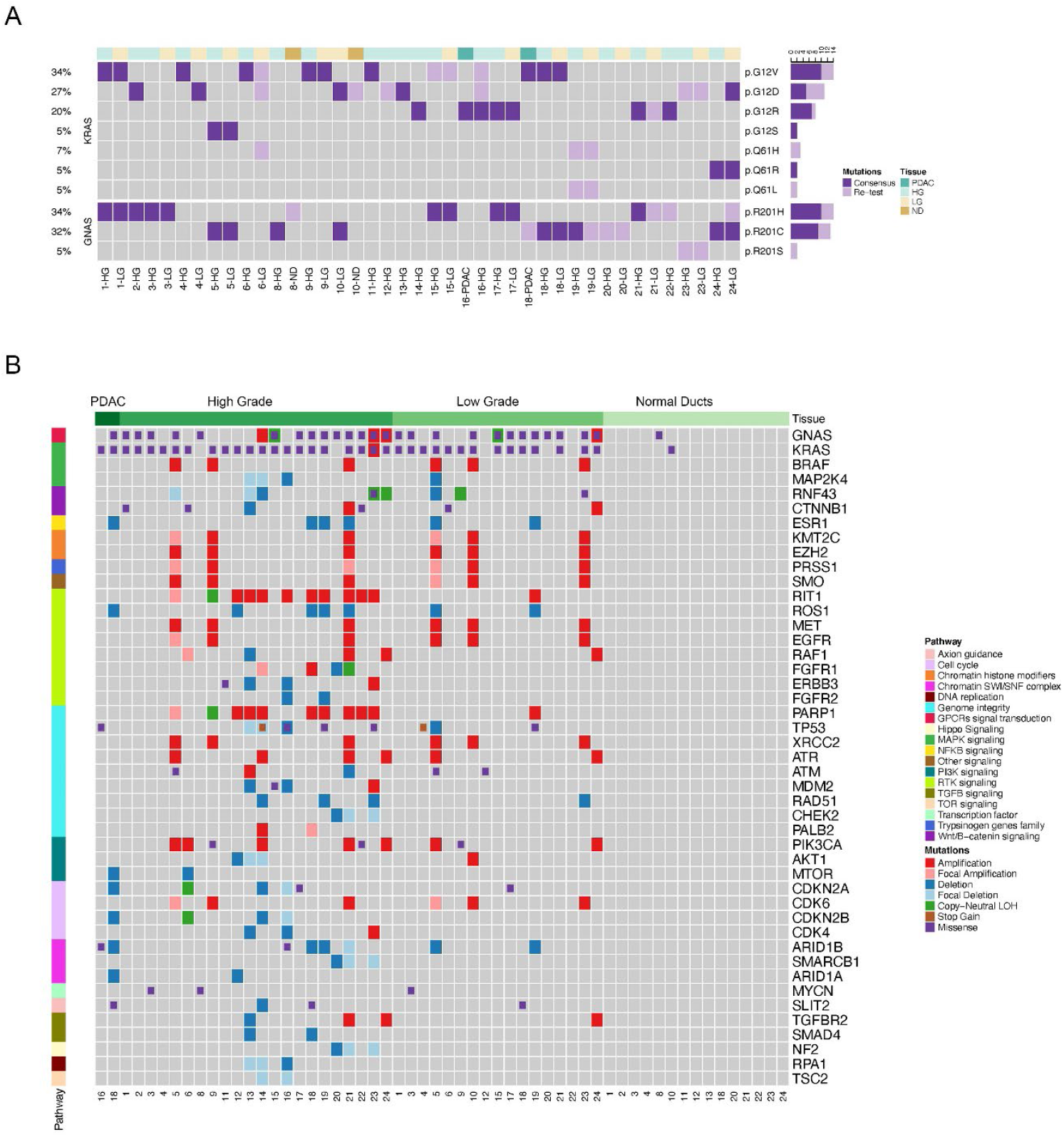
Genomic landscapes of IPMN lesions. **A)** Single nucleotide variants identified in *KRAS* and *GNAS* in LG and HG IPMNs. Dark purple: somatic mutations detected for *KRAS* and *GNAS*. Light purple: mutations detected by re-testing. **B)** SNVs and CNAs identified in ND, LG, HG and PDAC regions and classified by relevant PDAC-related pathways.

### Genetic alterations in premalignant lesions

On average, 34 non-synonymous mutations were detected per micro-dissected LG and HG region with a mean mutation burden (MB) of 0.91 mutations per megabase (Mb) which agrees with previous reports (41 mutations and 1.11 MB, (19)). We compared the mutational load from LG to HG lesions and found no differences in the number of mutations by an unpaired and paired analysis (**Supplementary Figure 2 A-B**). We analyzed CNA events in pre-cancer lesions and classified alterations as focal when smaller than 3 Mb (average 0.8 Mb and median 0.65 Mb). Those CNAs not classified as focal had an average size of 17Mb (median 8Mb). Mutations and CNAs were identified in genes belonging to pathways such as MAPK, RTK, and TGFB signaling, as well as genome instability and cell cycle (**Figure 1B)**. Among others, these CNAs involved genes in RTK signaling (*RIT1, ROS1, MET, EGFR, RAF1, FGFR, ERBB3, FGFR2*), genome integrity (*TP53, PARP1, XRCC2, ATR, ATM, MDM2, RAD51, CHEK2, PALB2*) and cell cycle (*CDKN2A, CDK6, CDKN2B, CDK4*) (**Figure 1B and Supplementary Figure 2C**). Overall, chromosomal alterations were more common than SNVs in IPMN lesions.

We then calculated an aneuploidy score (AS) for each sample as described previously, defined as the number of chromosome arms spanned by a CNA (minimum 75% of the chromosome arm) [23]. The AS was significantly higher in HG regions compared to LG regions (**Figure 2A - B**). One of the most frequent alterations was amplification of the 1q arm (**Supplementary Figure 2C**) (9 patients, 10 HG or LG regions). We performed fluorescence in situ hybridization (FISH) as an orthogonal validation method. Probes expanding locations 1q12, 1q21, 1q22, 1q telomere and 1p32 (control) were utilized in 2 cases containing paired LG and HG lesions (IDs 18 and 23); all q probes showed a significantly higher mean count field foci compared to normal ducts (**Figure 2C-D**). In addition, 1q amplification was more common in HG (9/23) *versus* LG (1/17) and the coding regions for poly (adenosine diphosphate [ADP] - ribose) polymerase (*PARP1*) and Ras Like Without CAAX 1 (*RIT1*) were located within the amplified loci. Thus, we performed integrated analysis by RNAseq and found significant overexpression of transcripts for both genes in lesions with gains in 1q compared to unaltered regions (**Supplementary Figure 2D-E**). Further, all IPMN lesions that showed 1q amplification showed a higher AS (p<0.0001) but not higher TMB (p>0.05) (**Supplementary Figure 2 F-G**). In this regard, *PARP1* plays a critical role in the DNA damage repair, including highly error prone DNA repair that enhances genomic instability [24, 25]. In addition to *PARP1*, we also investigated the presence of genetic alterations in other chromosomal instability related genes. 22 out of 39 pre-neoplastic lesions harbor SNVs and/or CNAs in *PARP1, TP53, XRCC2, ATR, ATM, MDM2, RAD51, CHEK2*, and *PALB2* which correlated with significantly higher AS in these lesions (**Supplementary Figure 2H**). In summary, CNAs appear to be common event in non-invasive IPMNs, increase upon progression to HG lesions, and specifically, 1q amplification appear to stratify for IPMNs at higher risk of progression.

**Figure 2:**
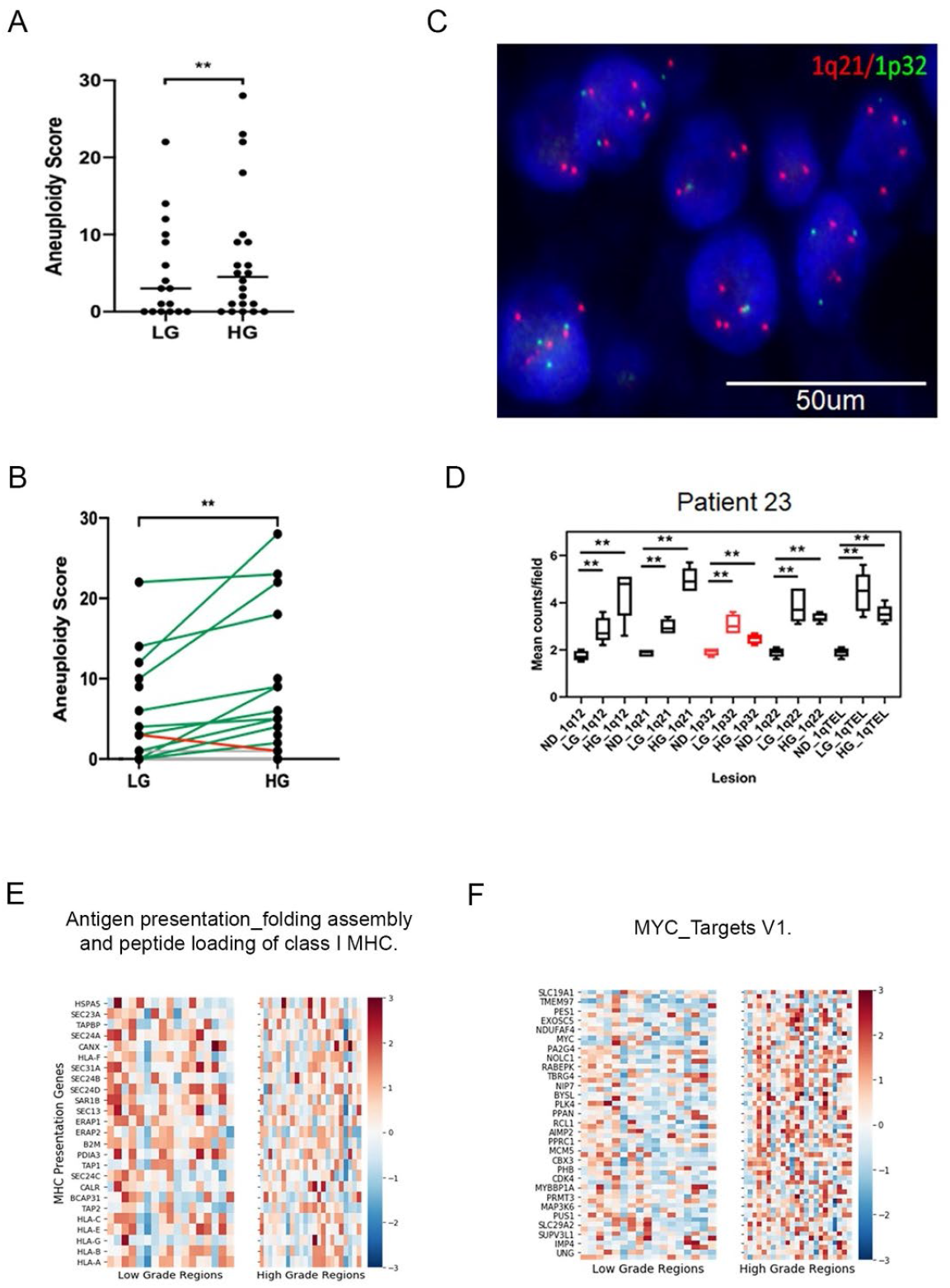
Comparison of CNAs between LG and HG IPMNs. Aneuploidy score between LG and HG by unpaired **A)** and paired **B)** analysis. Validation **C)** and quantification **D)** of chromosome 1q amplification by FISH. GSEA analysis of pathways enriched in LG **E)** and HG **F)** IPMNs.

### Transcriptomic analysis of IPMNs

In order to evaluate transcriptomics signatures driving the LG to HG progression, we analyzed bulk RNAseq data from microdissected LG and HG lesions that were processed by the SMART3 approach [14]. Differential gene expression followed by gene set enrichment analysis (GSEA) between HG and LG lesions indicated that LG lesions were enriched in pathways associated with antigen presentation compared to HG lesions (NES: −2.123, FDR: 0.0017, **Figure 2E**) while HG lesions were enriched for pathways more closely related with PDAC biology such as oncogenic MYC targets (NES: 2.657, FDR:0.0), E2F targets (NES: 2.772, FDR: 0.0), cell cycle and DNA replication (**Figure 2F** and **Supplementary Figure 3A**). In addition, we characterized the expression of potential neo-epitopes and their presentation. We applied WES data to OptiType [26] and PolySolver [27] to perform neo-epitope prediction and identified 20 genes with at least 10 potential neo-epitopes which were significantly enriched in HG compared to LG (**Supplementary Figure 3B-C**). Despite the putative higher neoantigen load, the downregulation of transcripts associated with the antigen presentation machinery in HG lesions suggests that immune evasion in cancer has its origins within the precursor microenvironment (PME) of non-invasive precursor lesions

Next, we studied whether known molecular PDAC subtypes could be identified in early premalignant HG and LG lesions. Using commonly accepted gene sets [28], 81% of all regions analyzed were classified into PDAC subtypes (**Supplementary Figure 3D-E)**. While in some IPMN samples more than one subtype was detected (Patient ID 2 and 20), in others, a class switch was found upon progression (**Supplementary Figure 3D)**. For example, the IPMN lesion in patient 20 experienced a subtype class switch (classical to basal) upon progression from LG to HG. Of note, acinar cells and normal ducts were almost exclusively classified in the ADEX/Exocrine group while the two PDAC samples were mostly basal (**Supplementary Figure 3D**). There was no difference between classical and basal assignment of IPMN lesions with regards to overall genomic characteristics like MB or markers of genomic instability like aneuploidy (*data not shown*). Nonetheless, basal lesions only harbored *KRAS* mutations while classical IPMNs contain mutations in both *KRAS* and *GNAS* (**Supplementary Figure 3F**).

### Evolutionary trajectory in pre-cancer lesions: SNVs *versus* CNAs

Previous work has demonstrated a highly heterogeneous progression pattern of IPMNs by SNVs, which is confirmed in our cohort [6, 7, 29, 30]. Clonal evolution was evaluated by two different approaches. First, clonal lineages were inferred using the Metastatic And Clonal History INtegrative Analysis (MACHINA), an algorithm that models the evolutionary trajectory and migration histories of clones in metastatic cancer using SNV data [31]. We categorized the SNV generated phylogenetic trees for each patient as “linear” or “branched”. Linear was defined when a clone in the LG acquired mutation(s) in a stepwise manner to give rise to a dominant clone present in the HG. Branched evolution was defined by early branching of the HG and LG lineages. SNV analysis indicated that 11/23 (48%, ID cases 1, 3-5, 9, 15, 17-19, 21 and 23) cases showed a linear evolution, while in the minority of cases, 5/23 (22%, ID cases 6, 12, 20, 22 and 24), the HG lesions showed a branched evolution from LG. In addition, in 7/23 (30%) the evolution could not be inferred. We then compared the molecular subtypes derived from RNAseq in lesions with their evolutionary trajectory based on SNV calls. In patients with linear evolution, the molecular subtype persists in most cases (9/11) during transition from LG to HG, while in patients with branched evolution, the molecular subtype of the LG IPMN is present in only 1/5 matched HG lesions (**Supplementary Figure 3D**).

To model the evolutionary trajectory of CNAs, we applied the Copy-Number Tree Mixture Deconvolution (CNTMD). This method uses multiple samples of a tumor and aims to build evolutionary trees [32]. We derived clonal evolution from CNAs by including events 5Mb and higher. Using this approach 5/23 (22%) HG lesions seem to follow a linear evolution by CNA analysis, 7/23 (30%) were branched while 11/23 (48%) could not be classified. In majority of cases (8/12) the evolutionary branch giving rise to the largest HG clone was associated with alterations in chromosome 1q (**Figure 3**). When comparing clonal evolution derived from CNAs and SNVs there was 42% (5/12) agreement. However, in 58% (7/12) the evolutionary trajectories were different and in four of these cases, it was due to CNA-driven branching evolution. These results indicate that evolutionary trajectories solely based on SNVs could poentially miss CNA-driven sub-clonal evolution and thus underestimate a hidden branching lineage that may facilitate IPMN progression. Overall, our work reiterates the previously described heterogeneity that characterizes evolutionary trajectories of IPMN progression, which is further accentuated with the consideration of CNAs.

**Figure 3:**
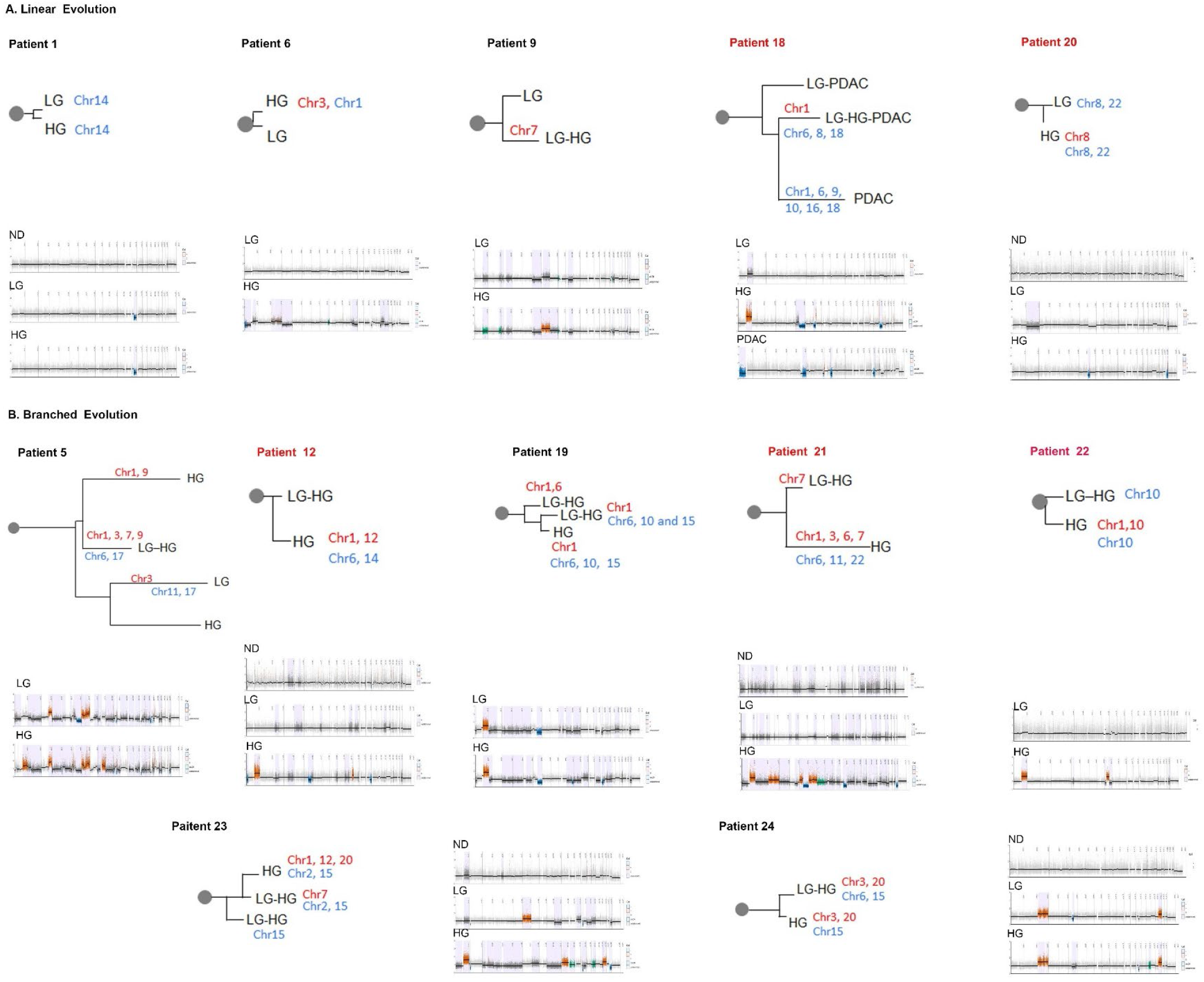
Inferred evolutionary trajectory derived from CNAs. Cases showing linear **A)** and branched **B)** evolution are depicted. For each case, evolutionary tree and segmentation plots are shown. Branches are drawn to scale based on the number of CNA events. Chromosomal aberrations associated to each branch are shown. Red indicate gains and blue losses. Patient ID highlighted in red indicate cases with coexisting PDAC.

We compared the evolutionary trajectory between cases with co-occurring carcinoma and those that did not progress to PDAC. In addition to the two cases in which we were able to analyze PDAC in the tissue blocks at hand (patient IDs 16 and 18), there were 7 additional samples with co-existing carcinoma (patient IDs 8, 9, 12, 14, 20, 21 and 22) in the final pathology report. At the DNA level, these cooccurring IPMN samples showed no differences in either MB or AS between LG or HG. Importantly, 7/9 showed chromosomal aberrations in chromosome 1q, five with amplifications, one with a partial amplification and an additional case, with a 1q LOH. Except for patient 8, the remaining cases with co-occurring carcinoma harbor mutations in chromosomal instability related genes, and in the cases with 1q amplification the *PARP1* locus was detected. This emphasizes the potential significance of a chromosome arm 1q amplification as a progression marker in IPMN.

To integrate MACHINA and CNTNB analysis and gain a better understanding of the evolutionary trajectory associated with IPMN progression, we analyzed 3 cases in detail. Patient 9 LG and HG lesions shared an ancestral lineage defined by *KRAS^G12^, PIK3CA^E542K^* and 66 additional SNVs. This clonal lineage gave rise to a dominant clone which contains 8 additional SNVs, present at 2% in the LG, expanding to become a major clone in the HG (**Figure 4A**). In agreement, the evolutionary trajectory derived from CNAs indicated a linear evolution for HG which was marked by an amplification in chromosome 7 (**Figure 4B and C**). In patient 21, LG and HG lesions shared a common ancestor carrying *KRAS^G12R^* and *GNAS^R201C^* mutations. The evolutionary phylogeny derived from SNVs indicates linear evolution of LG to HG where in the latter, 14 additional mutations are acquired (**Figure 4D**). In contrast, the inferred clonal evolution derived from CNAs show evidence of CNA-driven branching with two clones present in the HG lesions. A major clone, unique for the HG lesion, showed gains in chromosomes 1, 3, 6 and 7 and losses in chromosomes 6, 11 and 22 which was confirmed by segmentation analysis (**Figure 4D-F)**. In addition, a second clone shared between the LG and HG was exclusively characterized by chromosome 7 amplification. For patient 18, the SNV data showed clones in LG, HG, PDAC as having a most recent common ancestor clone with truncal *KRAS^G12V^* and *GNAS^R201C^* mutations. While there is a sub-clonal population unique to the LG and HG, the PDAC evolved independently from the dominant clone present in HG and LG (**Supplementary Figure 4A**). Instead, a minor subpopulation, present at 3% in the HG, later expanded giving rise to the dominant PDAC clone defined by an *MTA1* mutation. Similarly, CNA-derived evolution showed a linear evolution from LG to HG which was characterized by gain in chromosome 1 and losses on chromosomes 6, 8, 18 as it was later clearly confirmed by chromosomal segmentation analysis. Of note, although both HG and tumor showed loss on chromosome 6, upon close inspection these losses occurred on different chromosomes i.e. one lineage lost the maternal copy while the other lost the paternal copy. In addition, two additional branches were detected, the first branch was shared between the LG and PDAC while in the second branch, a completely independent PDAC clone, was identified with losses on chromosomes 1, 6, 9, 10, 16 and 18 (**Supplementary Figure 4B-C**).

**Figure 4:**
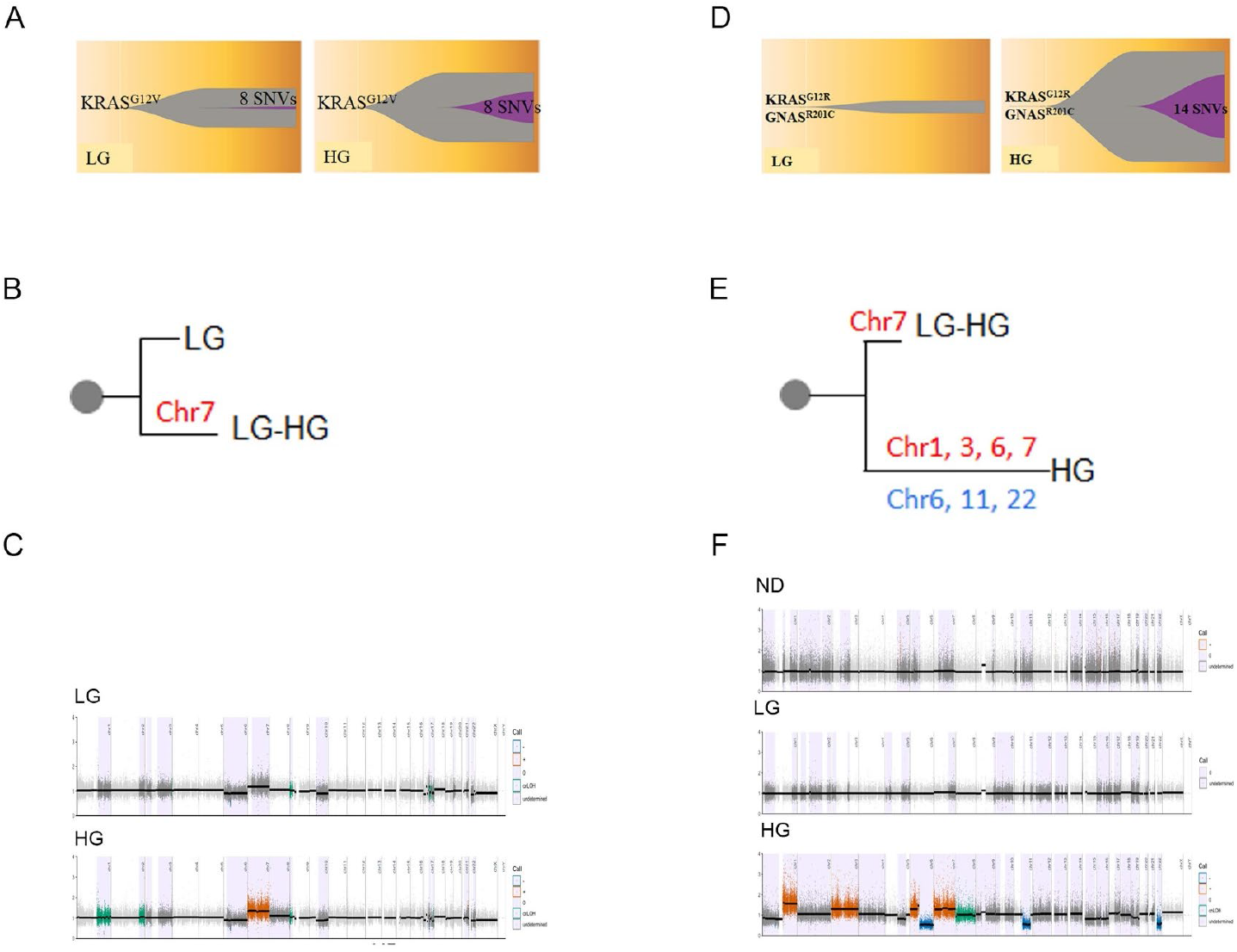
Evolutionary trajectory of IPMN lesions. A case with linear **A-C)** and branched **D-F)** evolution is depicted. **A** and **D)** Muller plots showing SNV-derived evolution for patients 9 and 21 respectively. **B** and **E)** CNA-derived clonal evolution showing main CNA events (red for gains and blue for losses). **C** and **F)** Segmentation plots intersected with HapLOHseq calls (lavender background).

## Discussion

In this multi-institutional study, we interrogated the evolutionary trajectories and transcriptomic aberrations that occur during IPMN progression, using paired whole exome DNA sequencing and whole transcriptome sequencing. The majority of IPMNs within our dataset harbor somatic “hotspot” mutations in *KRAS* and *GNAS* as an early event, consistent with previously reported findings [33]. In contrast to the prior study by Fischer and colleagues that found striking heterogeneity in driver gene mutations in IPMNs [30], we found driver SNVs to be relatively homogeneous, which may be the consequence of our more limited multi-region sequencing. On the contrary, our study suggests CNA events play a more pervasive role in the IPMN progression models than previously appreciated, such that accumulation of CNA events in a subgroup of LG IPMN seem to pave the way to further progression. In particular, we demonstrate that HG samples with co-occurring PDAC tend to frequently harbor chromosomal 1q amplifications. Chromosome 1q amplification has also been detected in precancer lesions in other cancer types such as breast [34] and esophageal [35]. In the context of PDAC, chromosome 1q amplifications have been previously described through single nucleotide polymorphism (SNP) arrays and microarray-based comparative genomic hybridization (CGH), but their relevance in non-invasive precursors is relatively unknown [36, 37]. The amplified region of chromosome 1q in IPMN harbors *PARP1*, whose product is an enzyme pivotal to DNA damage repair, including homologous recombination and error-prone DNA repair processes. An increase in PARP-1 enzymatic activity has been associated with the highly error-prone DNA repair pathway known as microhomology-mediated end joining (MMEJ), which has been reported to increase chromosomal structural alterations and genomic instability [38, 39]. In our data, amplification in chromosome 1q, with concomitant overexpression of *PARP1*, was also associated with an increase in AS, suggesting that this event might be a prelude to genomic instability-enhanced progression in IPMNs.

We also confirm previous findings that from an evolutionary standpoint, IPMN progression is quite heterogenous [6, 7, 29, 30]. Evolutionary modeling of our datasets demonstrates that while both linear and branched trajectories are present, a majority of IPMNs with co-occurring invasive phenotypes follow a branched evolution. Previous data had supported this evolutionary track with multi-region analysis of SNVs [7], but we now demonstrate how CNA-based phylogeny (with the added integration of transcriptome data) can identify similar patterns even with more limited sequencing analyses. For example, in patient 18 we found that the HG and PDAC areas show two distinct parental chromosomal deletions in chromosome 6q, distinct alterations in chromosome arms 1p and 1q and a transcriptomic class switch on RNA sequencing, all of which point towards an independent development of the invasive PDAC from LG evolving in parallel to a co-occurrent HG lesion. Unfortunately, our results also indicate the challenges of predicting the pattern of progression in any given LG IPMN, given the inherent heterogeneity of possible pathways to HG and beyond.

Finally, it is worth noting that established transcriptomic PDAC signatures [17–19, 28] are also detectable in nearly all IPMN lesions. Signatures that have been previously suspected as likely originating from non-neoplastic tissue (exocrine-like and ADEX) were almost exclusively seen in acinar cells and normal ducts [18, 19], whereas LG, HG and PDAC were mostly classified into the basal/squamous or classical/immunogenic subtypes. This demonstrates that even premalignant lesions express both consensus PDAC signatures and that certain pathways attributed to invasive carcinomas are even present within these lesions. The transcriptomic data also revealed that reduction in transcripts associated with MHC class I antigen presentation machinery was a feature of HG IPMNs. Recent work has shown reduced expression of MHC-I at the cell surface of PDAC cells which are targeted for lysosomal degradation [40, 41]. Our data confirms that perturbation of antigen presentation might occur even in non-invasive precursor lesions, and that there are other mechanisms beyond protein recycling to the lysosomes that might contribute to this dysfunction. Recent immune profiling data of IPMN progression has shown that the PME of non-invasive IPMNs is altered towards a more immune suppressive milieu upon progression to HG, and remarkably, comparable immune alterations are also observed in the matched LG IPMNs prior to progression [42]. Thus, the PME plays an integral permissive role in IPMN progression, with likely multiple mechanisms through which effective antigen presentation is perturbed early in multistep neoplasia.

## Acknowledgments

We thank the patients and their families and all MCL Pre-Cancer Atlas Pilot Collaborative Project centers for providing samples and critical feedback. The authors thank Mr. Bret M. Stephens and Dr. Mark Hurd from MD Anderson for sample processing and acquisition, Dr. Kristen Jepsen, and Mrs. Huazhen Yao at the UCSD Institute for Genomic Medicine Genomic Center for sample processing, and Drs. Hidetoshi Mori and Alexander D. Borowsky from the University of California Davis for providing helpful feedback.

A. Maitra is supported by the MD Anderson Pancreatic Cancer Moon Shot Program, the Khalifa Bin Zayed Al-Nahyan Foundation, and the NIH (U01CA196403, U01CA200468, and P50CA221707). V. Bernard is supported by the NIH (U54CA096300, U54CA096297, and T32CA217789). J. J. Lee is supported by the NIH (T32CA009599). A. Semaan is supported by the German Research Foundation (SE-2616/2–1). J. Wong is supported by T32 CA217789. S. Sen is supported by UO1CA 214263. M. Yip-Schneider and M. Schmidt are supported by grants U01CA196403, U01CA200468. Y Jakubek is supported by NCI K22CA258678. A. Lowy is supported by NIH R21CA273974-01. Support for collection of University of Utah samples used in this study was partially provided by subcontracts of A. Maitra’s listed grants (U01CA196403, U01CA200468) as well as P30CA042014 to the Huntsman Cancer Institute for support of core facilities. D. Nachmanson is supported by California Tobacco Related Disease Research Program pre-doctoral fellowship to DN (28DT-0011). O. Harismendy is supported by U01CA196406, U01CA196406-03S1, T32GM008806, T15LM011271

## Data availability

All datasets generated in this study have been uploaded to MCL-JPL LabCAS repository (https://mcl-labcas.jpl.nasa.gov/labcas-ui/m/index.html) and are available upon request.

## Figure and Tables Legends

**Supplementary Figure 1**: Schematic representation of the strategy employed. **A)** Correlation between block age and DNA integrity. **B)** Orthogonal validation of *GNAS* and *KRAS* mutations WES calls by ddPCR. **C)** Tissue sample showing areas isolated by LCM. **D)** Agilent’s chemistry was used to perform WES while Smart3’Seq was used for a whole transcriptomics approach.

**Supplementary Figure 2.** Comparison of SNVs and CNAs in LG and HG IPMNs. Mutational load between LG and HG by unpaired **A)** and paired **B)** analysis. **C)** Representation of CNA events detected in this study, red: gains, blue: losses, green LOH, gray: undetermined. Expression of PARP1 **D)** and RIT1 **E)** in cases with and without 1q amplification. Aneuploidy score **F)** and mutational burden **G)** for lesions with and without 1q amplification. **H)** Comparison of aneuploidy score in precancer lesions harboring SNVs or CNVs events in chromosomal instability (CI) related genes.

**Supplementary Figure 3.** Transcriptomic profile of IPMNs. **A)** GSEA analysis of pathways enriched HG lesions. **B)** Top 20 genes with 10 or more neoepitopes detected in this study. **C)** Analysis of neoepitopes from top 20 genes present in cases with paired LG and HG lesions. **D)** Expression of PDAC molecular subtypes in pre-cancer cystic lesions; 2-5 replicates were sequenced per region with each row representing classification by Moffit (top), Collisson (middle) and Bailey (bottom). **E)** FDR for Moffit, Collison and Bailey. **F)** Correlation between *KRAS* and *GNAS* mutation status and PDAC molecular subtypes.

**Supplementary Figure 4**. Evolutionary trajectory of an IPMN lesion with co-occurring PDAC. **A)** Muller plots depicting SNV-derived evolution for patient ID 18. **B)** CNA-derived clonal evolution showing main CNA events (red for gains and blue for losses). **C)** Segmentation plots intersected with HapLOHseq calls (lavender background)

**Supplementary Table 1**: Comparison in MAF and sequencing depth between a targeted and WES approach.

