## Supplementary figures and images for "Integrated Molecular Characterization of Intraductal Papillary Mucinous Neoplasms: An NCI Cancer Moonshot Precancer Atlas Pilot Project"

### Supplementary Figures

Supplementary Figure 1

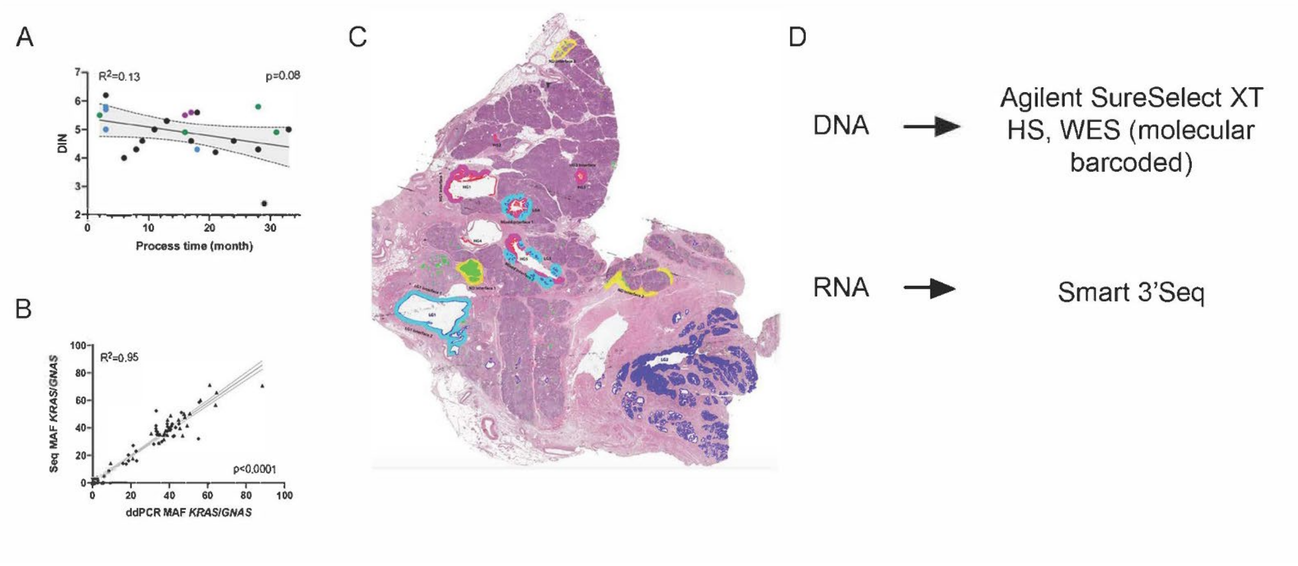

Supplementary Figure 2

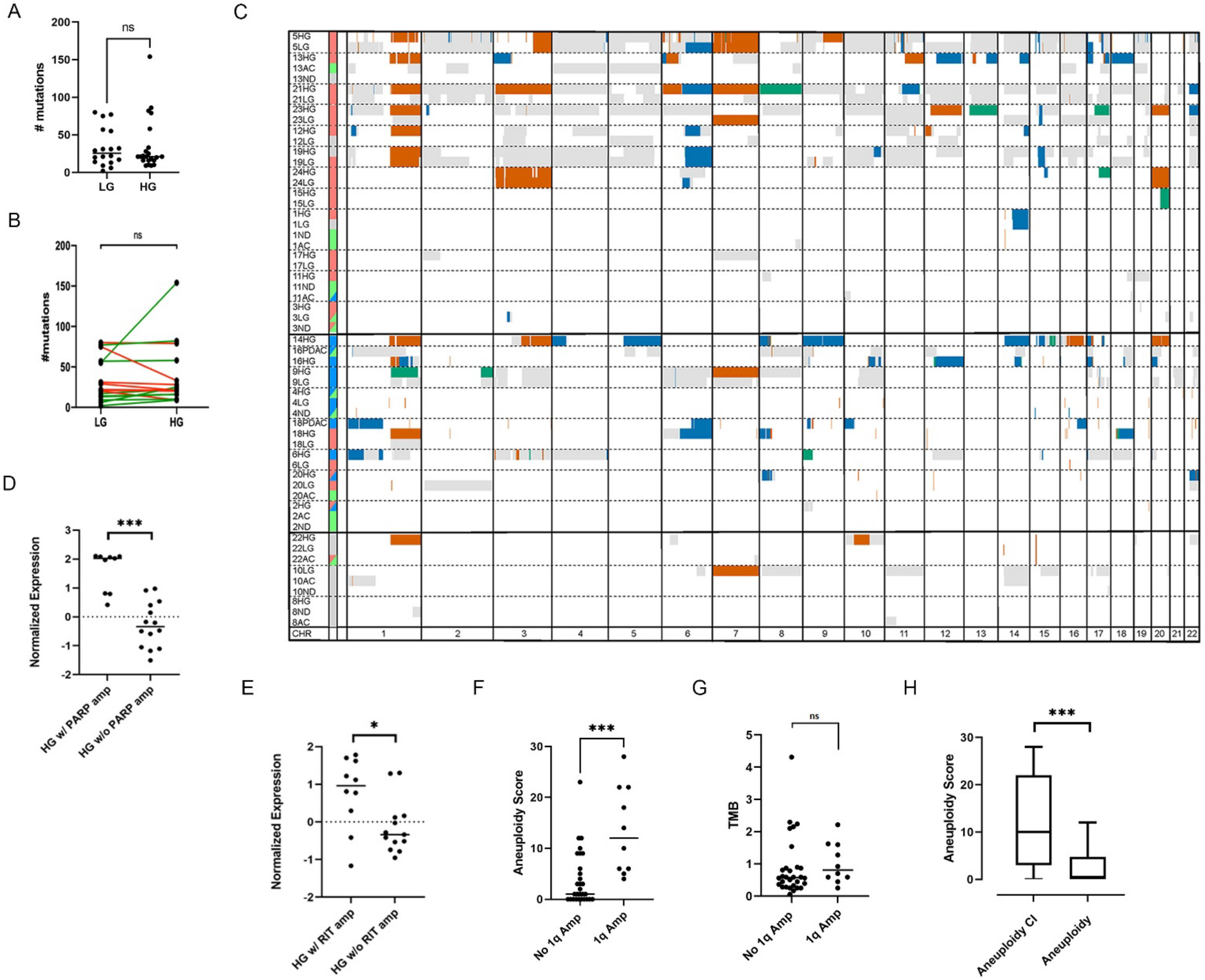

#### Supplementary Figure 3

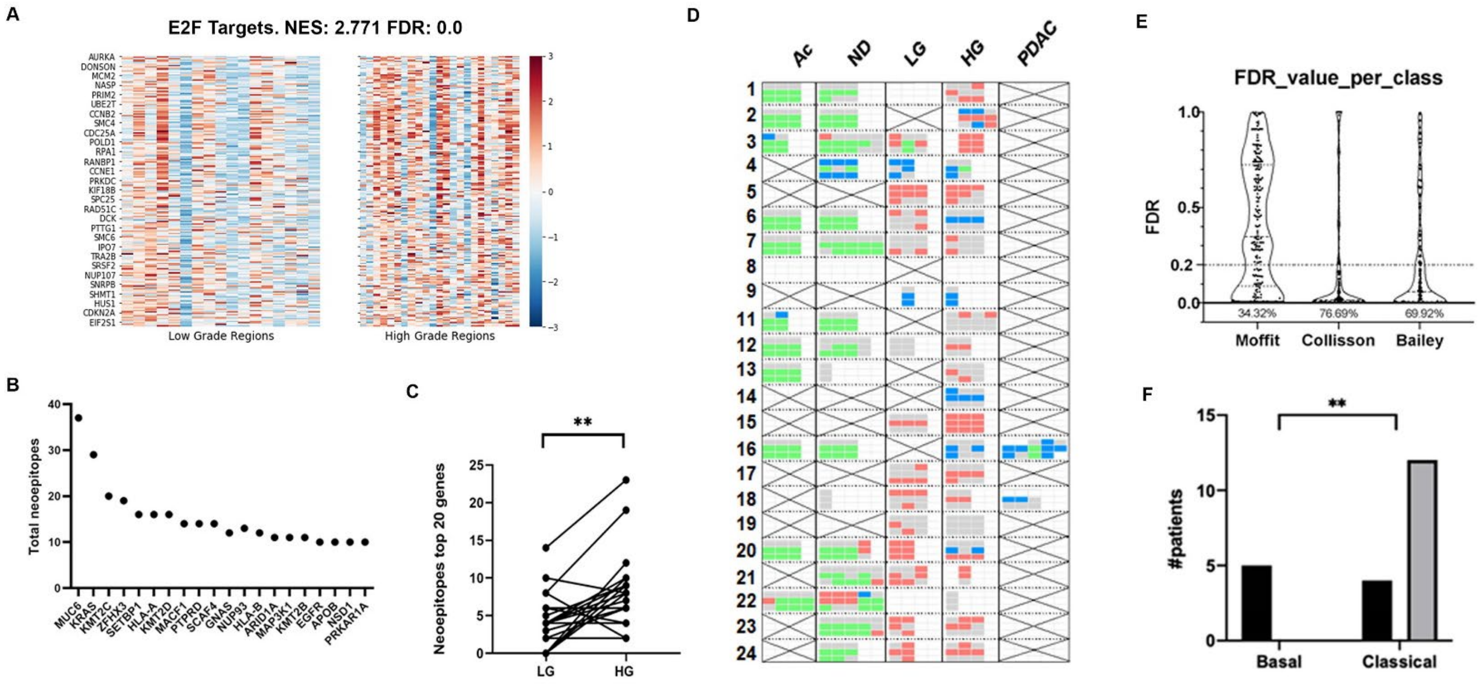

#### Supplementary Figure 4

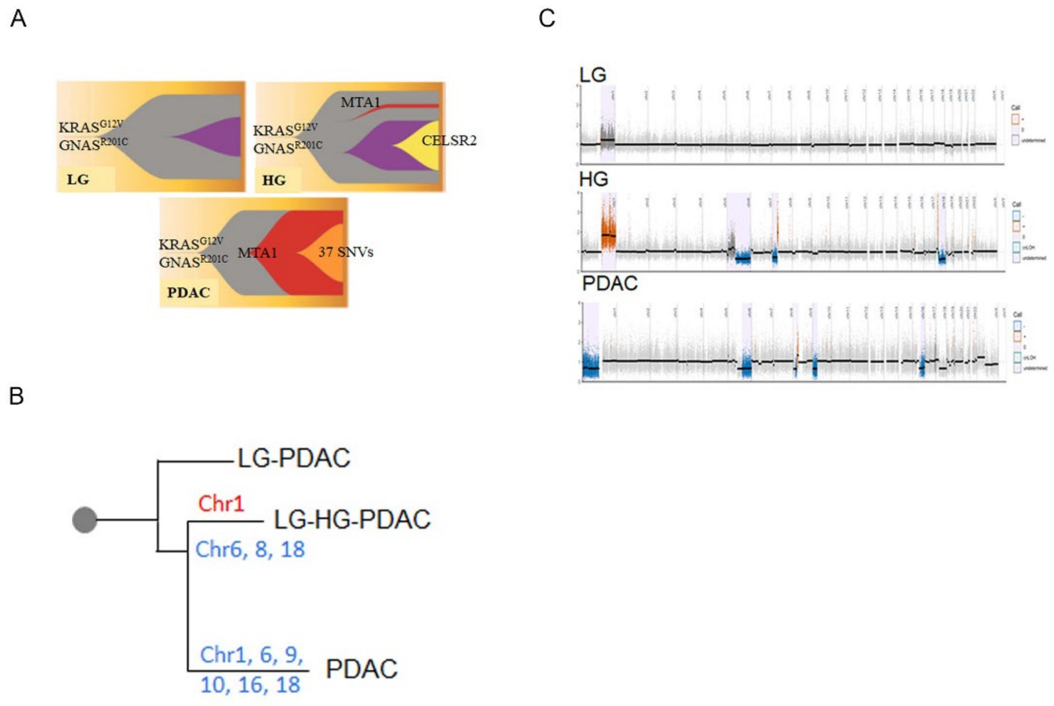
